# Co-cultivation with Azolla affects the metabolome of whole rice plant beyond canonical inorganic nitrogen fertilization

**DOI:** 10.1101/2024.10.02.615589

**Authors:** Elena Consorti, Alma Costarelli, Sara Cannavò, Martina Cerri, Maria Cristina Valeri, Lara Reale, Antonietta Saccomano, Chiara Paleni, Veronica Gregis, Martin M. Kater, Federico Brilli, Francesco Paolocci, Andrea Ghirardo

## Abstract

Azolla spp. are floating ferns used for centuries as biofertilizers to enrich the soil with inorganic nitrogen and improve rice yields. In this study, rice plants were grown together with Azolla by maintaining a low and constant concentration of inorganic nitrogen. We employed a combination of non-targeted metabolomics, chemometrics, and molecular networking to dissect the impact of Azolla co-cultivation on the metabolome of rice roots-and leaves. Our analyses revealed that Azolla releases a broad range of metabolites in the culture medium, mainly comprising small peptides and flavonoids. Moreover, in rice co-cultivated with Azolla, we observed a systematic response in the upregulation of metabolites that started from the roots and, over time, shifted to the leaves. During the early stages of co-cultivation, Azolla led to the accumulation of small peptides, lipids, and carbohydrates in roots, and flavonoid glycosides and carbohydrates in leaves of rice. Consistent with these results, transcriptomics analysis of rice roots indicated significant changes in the expression of genes coding for small peptide and lipid transporters, and genes involved in amino acid salvage and biosynthesis. Overall, our study highlights novel growth-promoting effects of Azolla on rice which could facilitate the development of sustainable techniques to increase yields.

**Highlights:** The aquatic fern Azolla synthesizes and releases a broad range of growth promoting metabolites (i.e. small peptides) that can be absorbed by the roots of co-cultivated rice plants

## Introduction

The major challenge of the twenty-first century is to sustainably feed a world population expected to reach ∼9.7 billion by 2050 (United Nations, 2019). Climate change (Das Gupta, 2014) and concerns about using chemical fertilizers and pesticides call for innovative strategies to achieve increased yields while decreasing the environmental impact of global crop cultivation (Matson et al., 1997). A strategy for the transition to a more sustainable agricultural production relies on the co-cultivation of crops with companion plants and associated microbes that fix atmospheric nitrogen, thereby acting as soil biofertilizers. An excellent example is the use of the *Anabaena* (*Trichormus*)*-azollae* symbiosis system as a sustainable source of nitrogen in rice cultivation (Watanabe and Liu, 1992). Azolla spp. is a small floating fern (Lumpkin & Plucknett, 1980) whose leaflets have cavities that provide a microenvironment for the nitrogen-fixing filamentous cyanobacterium *Anabaena* (*Trichormus*) *azollae* (Kumar et al., 2019). *Azolla-Trichormus*, hereinafter referred to as “Azolla”, is a unique symbiotic system that persists throughout the fern’s life cycle and allows to double its mass in 3-5 days. The Azolla nitrogen-fixing capacity, accounting for 30-40 kg N ha^-1^ in two weeks when growing in nitrogen-free solution (Watanabe et al., 1977), is higher than that achieved by the symbiosis between legumes and Rhizobia and enables the fern to grow in waterlogged habitats poor in nitrogen content (Bhuvaneshwari et al., 2015). Given the high growth rate and great N-fixing potentials, Azolla can cover large water basins in a very short time and further enrich the soil with nitrogen which is slowly released after plant death and decomposition (Mahanty et al., 2017). The inorganic nitrogen released by Azolla and available to the companion crops, such as rice, is about 70% of that of ammonium sulfate (Watanabe et al., 1977). In Indian paddy soils, Azolla decomposed in 8-10 days to benefit to the co-cultivated rice after 20-30 days (Singh 1977). For this reason, Azolla has been used for centuries as biofertilizer in rice paddies in China and Vietnam (Singh, 1989; Watanabe, 1982; Watanabe et al., 1989, Bhuvaneshwari et. al, 2012, 2013; van Hove and Lejeune, 2002), and it is still currently used either as green manure or intercropped with rice (Okonji et al., 2012). Thus, the role of Azolla in supplying inorganic nitrogen to rice fields is well documented (Peters and Meeks, 1989). In addition, it has been demonstrated that Azolla, following decomposition, increases soil mineral content (N, P, K, Ca, Mg, and Na) and organic matter (Bhuvaneswari et al., 2013).

Studies on free-living extracts of Azolla have indicated that this fern has the potential to produce hormones, vitamins, and other growth-promoting substances that enhance crop growth (Misra & Kaushik, 1989a, 1989b; Wang et al., 1991; Mofiz et al. 2023). Moreover, the growth-promoting effect of Azolla starts early, before the end of its life cycle, suggesting that molecules stimulating plant growth are released by the Azolla into the surrounding environment while it is still alive. Evidence has shown an increase in rice plant height and number of tillers in rice following addition of Azolla to the soil (Bhuvaneshwari et al., 2015). It has also been shown that co-cultivation of rice with Azolla and associated cyanobacteria boosts rice growth at an early stage by increasing root and shoot growth and, ultimately, enhancing the rice grain weight and protein content (Venkataraman and Neelakantan 1967; Singh and Trehan 1973 Bhuvaneshwari, 2012). Thus it has been postulated that Azolla may be a source of growth-promoting compounds released into the water (Wagner 1997), although no report has been published to confirm it. However, the molecular mechanisms by which Azolla exerts its growth-promoting effects on co-cultivated crops remain unclear, and farmers still prefer to rely on chemical fertilizers to control rice yield (Marzouk et al., 2023).

Moving from the companion study by Cannavò et al. (2024, Preprint), which demonstrated the morphological and transcriptional changes in rice induced by co-cultivation with Azolla, here we employed non-targeted metabolomics to investigate the alterations in the rice plant metabolome triggered by the fern. Specifically, our objectives were to: i) study the Azolla-induced changes of the metabolomes in leaves and roots of rice plants at two time points following the onset of Azolla-rice co-cultivation; ii) to identify the metabolites released by Azolla into the growth medium and evaluate their potential role in influencing rice phenotype and growth.

## Materials and methods

### Plant material and experimental setup

*Azolla filiculoides* Lam. employed in this study were collected, characterized, and grown under controlled conditions in Watanabe solution (Table S1; Watanabe et al., 1992) as reported in Costarelli et al., (2021). To set the co-cultivation experiments rice (*Oryza sativa* cv. Kitaake) seeds were sterilized and germinated in Petri dish as reported in Cannavò et al. (2024, Preprint). Rice plants were grown hydroponically in Yoshida solution (Table S2; Yoshida et al., 1976) by employing expanded clay balls (Atami, Netherlands) as plant support, and grown under controlled environmental conditions in 50x33x11 cm (length/width/depth) boxes filled with 6 L solution and placed in a climatic chamber with a temperature of 25/20°C (day/night), photosynthetic photon flux density (PPFD) of 220 µmoles m^-2^ s^-1^ provided by fluorescent tubes (Philips, Netherlands) and a 10-h photoperiod.

A set of 6 boxes were prepared to grow 4 rice plants each in Yoshida solution: 3 boxes containing a total of 12 rice plants were co-cultivated with Azolla (+AZ), and 3 boxes containing other 12 rice plants that were cultivated without Azolla (-AZ) (Fig. 1a). All the boxes were wrapped and darkened with aluminum foil to prevent the development of algae. The pH of Yoshida solution was adjusted with NaOH 1M to pH=5.0 and completely replaced every 2 weeks. Leaves and roots of (+AZ) and (-AZ) rice plants were sampled 40 and 60 days from the onset of hydroponic co-cultivation (doc). The roots and leaves were sampled from three different rice plants from each of the 6 boxes. All samples were flash-frozen in liquid nitrogen, freeze-dried, and stored (at 4°C) for non-targeted metabolomics analysis. To investigate the metabolites exchanged between rice and Azolla since the early phase of their interaction but when rice plants were already well acclimated to the hydroponic condition, the liquid culture medium was sampled 15 days from the onset of hydroponic cultivation, by collecting 10 ml from each (+AZ) and (-AZ) boxes. This 15-day time point coincides with the morphological and molecular analyses performed on the same rice plants in the companion study (Cannavò et al. 2024, Preprint). In addition, 10 mL of liquid culture medium was collected from the 3 boxes where Azolla was grown alone on Watanabe solution, under the environmental conditions described above, and from fresh Watanabe solution as control, and dried with speed vac (Thermo-Fisher scientific, USA).

**Figure 1.**
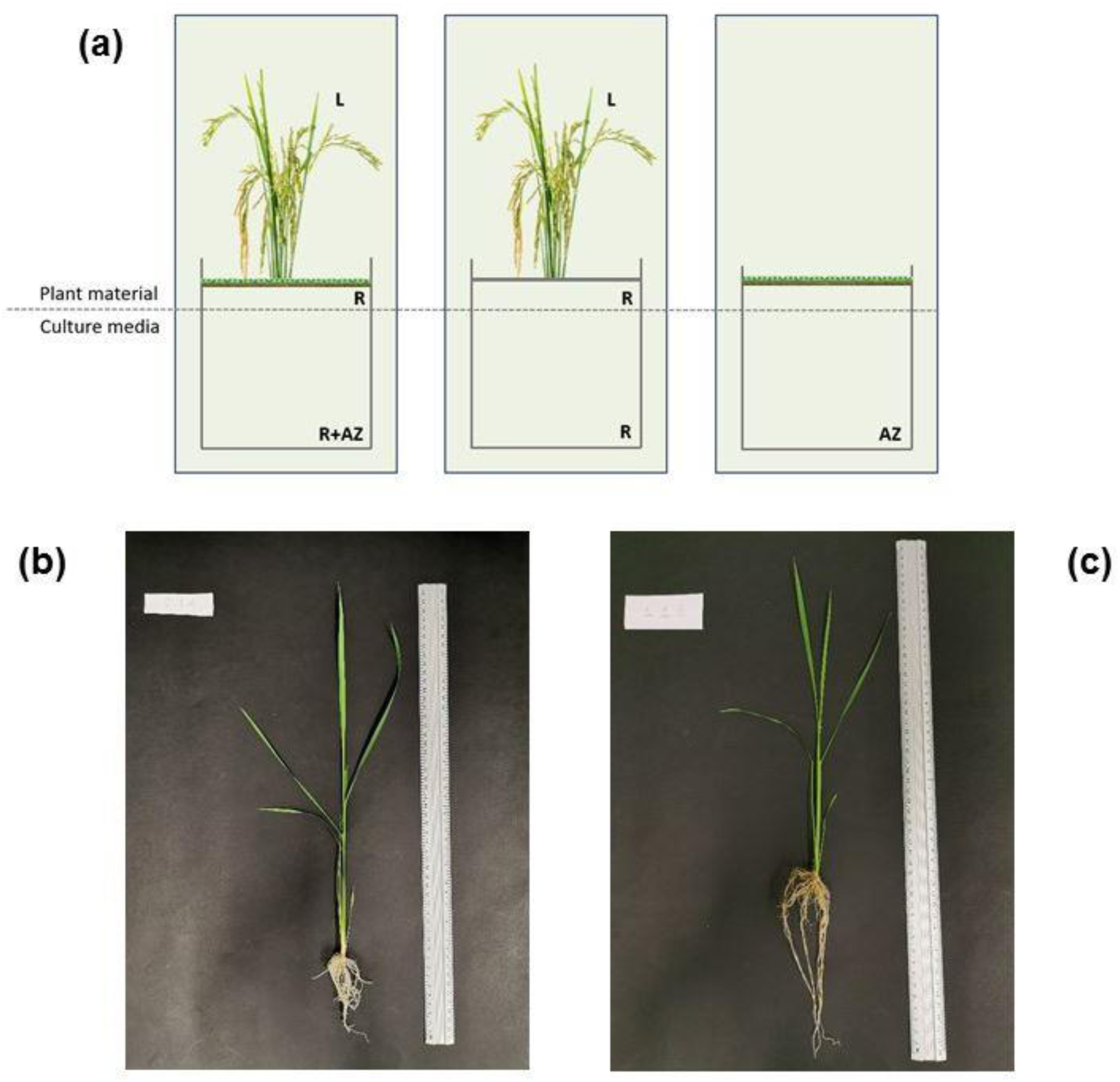
Schematic diagram of the experimental design (a); pictures showing the phenotypic differences of rice plants after 30 days of co-cultivation with-(b) and without-Azolla (c).

### Non-targeted metabolomics analysis by UPLC-UHR-QqToF-MS

The extraction of metabolites followed the protocol described in Bertić et al. (2021). Homogenized and powdered (+AZ) and (-AZ) rice leaf (L), root (R) samples, and lyophilized culture media samples were extracted with cold methanol:2-propanol:H_2_O (1:1:1, v/v/v) solution containing 50 µL L^-1^ of internal standard mixture (Table S3). The chemicals (LC-MS hyper grade) methanol/H_2_O were purchased from Merck (Darmstadt, Germany) and 2-propanol/acetonitrile from Honeywell (Puchheim, Germany). Due to plant material limitations, we extracted 25 mg of rice leaf with 1000 µL of solvent and 12.5 mg of rice root with 500 µL of solvent, i.e., using the same material-to-solvent ratio. Samples were mixed for 1 min inside a 2 mL polypropylene tube and sonicated in an ultrasonic bath for 10 min at 5°C. The solution was then centrifuged for 10 min at 10,000 rpm at 5°C. Four-fifths of the initial extraction volume was recovered and dried by SpeedVac (Univapo 150H, Uniequip, Planegg, Germany). The residue was dissolved in 350 µl of 50% (v/v) acetonitrile in water, mixed for 1 min, centrifuged for 10 min at 10,000 rpm at 5°C and the supernatant was ready for metabolic analysis. Culture media samples were directly dissolved in acetonitrile/water.

We strictly followed our established non-targeted metabolomics analysis (Ghirardo et al., 2020; Bertić et al., 2021) based on measurements with Ultra Performance Liquid Chromatography (UPLC) Ultra High resolution (UHR) tandem quadrupole/Time-of-Flight (QqToF) Mass Spectrometry (MS). The LC-MS instrument is composed of an Ultimate 3000RS UPLC (Thermo Fisher, Bremen, Germany), a Bruker Impact II (QqToF) and an Apollo II ESI source (Bruker Daltonic, Bremen, Germany). Each sample was measured twice, both on a reversed-phase liquid chromatography (RPLC) column and on a hydrophilic interaction liquid chromatography (HILIC) column (Bertić et al., 2021) to obtain an optimal separation of nonpolar and polar metabolites, respectively (Saba et al., 2001). We analyzed each sample with both RPLC and HILIC columns with MS operated both in positive and negative electrospray ionization modes (for details on chromatography and MS parameters, see Bertić et al., 2021). Data analysis followed Bertić et al. (2021). In short, raw data obtained from LC-MS were manually checked using the software Compass® Data Analysis 4.2 (Bruker Daltonik) for quality control, and corrupted chromatograms were discarded from the analysis. Data were further processed using Metaboscape 4.0 (Bruker) to perform isotope filtering, mass calibration, peak peaking, alignments, and peak-groupings based on peak-area correlation. Sample groups (i.e., +AZ and –AZ treatment, 40 and 60 duration of the treatment, L and R plant organ) were created in Metaboscape and only mass-features with >60 % presence at least in one group were retained for analysis. Intensity threshold and recursive counts were defined in Compass® and details on processing parameters can be found in Table S4.

Metabolite annotation was achieved by library comparison (Bertić et al., 2021), and we reported the non-annotated and non-classified metabolites as mass-features (MFs), giving the measured mass-to-charge ratio (*m/z*). For those MFs that were not found in databases, we used the recently developed multi-dimensional stoichiometric compound classification (MSCC) method, which classifies compounds based on their elemental composition in the chemical categories of proteins-related, amino sugars, lipids, carbohydrates, secondary metabolites (Rivas et al., 2018). Elemental composition of MF was calculated based on the exact measured mass and the sum formula was computed by the ‘SmartFormula function’ of Metaboscape. Molecular formulas were further used to calculate H:C, O:C, C: N, C:P, S:C, N:P ratios to depict Van Krevelen diagrams. It should be noted that multiple MF may relate to a single metabolite. Moreover, based on the chemical formula, metabolites were tentatively annotated by using the PubMed open database (https://pubchem.ncbi.nlm.nih.gov/) and National Institute of Standards and Technology (NIST) Chemistry WebBook, SRD 69 (https://doi.org/10.18434/T4D303). Systematic classification of tentatively annotated compounds and unknown metabolites were achieved by SIRIUS4 (Dührkop et al., 2019; 2020) using the tools CANOPUS (Djoumbou et al., 2016), CSI:FingerID and COSMIC (Dührkop et al., 2015; Kim et al. 2021; Hoffmann et al., 2022) on molecules that possessed fragmentation spectra (MS/MS) and were found to be statistically significant (adj. p-value < 0.05) between the comparison groups.

Molecular networks were created using the online workflow (https://ccms-ucsd.github.io/GNPSDocumentation/) on the GNPS website (http://gnps.ucsd.edu) (Wang et al., 2016). The data were filtered by removing all MS^2^ fragment ions within ± 10 Da of the *m/z* of the precursor. MS^2^ spectra were window-filtered by choosing only the first 6 fragment ions in the ± 50 Da window across the spectrum. The precursor ion mass tolerance was set to 0.05 Da and the MS^2^ fragment ion tolerance to 0.05 Da. Networks were then created in which edges were filtered to have a cosine score greater than 0.70 and more than 6 corresponding peaks. Furthermore, edges between two nodes were retained in the network when each of the nodes appeared in the respective top 10 most similar nodes. Finally, the maximum size of a molecular family was set to 100 and the lowest scoring edges were removed from the molecular families until the molecular family size was below this threshold. The network spectra were then searched in the GNPS spectral libraries. The library spectra were filtered in the same way as the input data. All matches maintained between the network and library spectra had to have a score above 0.7 and at least 6 matching peaks.

### RNA sequencing

For RNA isolation, three pools of rice root tips (up to 1 cm from apices) were collected 15 days after the onset of hydroponic cultivation from both control and Azolla co-cultivated rice plants. RNA isolation, cDNA library preparation, and RNAseq analyses were conducted as reported in Cannavò et al. (2024, Preprint).

### Statistics

All analyses of metabolomic data were performed on 3-6 independent replicates. Multivariate Data Analysis (MDA) was performed according to Bertić et al., (2021) and by using the software SIMCA-P v13.0.3.0 (Umetrics, Umeå, Sweden). Prior to analysis, data were always centered, transformed logarithmically (log10) and Pareto scaled (Eriksson, 1999; van den Berg et al., 2006). Orthogonal Partial Least Squares Discriminant Analysis (OPLS-DA) models were calculated using as Y-variables the rice plant treatment (+AZ and -AZ) and (40 and 60 doc) duration of the treatment (excluding plant material as Y) and assigning a binary discriminating variable codex to their class (plant organ, treatment and days of treatment). Once the model was created, it was auto-fitted by SIMCA® to the maximum number of significant components, the mass-features having a Variance Importance of Prediction (VIP) value > 2 were selected and further analyzed. All MF that had an abundance not statistically different from blanks were removed from the analysis. Significance was tested by *t*-test after correction for multiple tests with the Benjamini-Hochberg false discovery rate procedure (Benjamini-Hochberg, 1995; Glen, 2015). Only mass-features with adj. p-values <0.05 were considered in the result section. Significant perturbations in the metabolome were evaluated with hypergeometric tests, using the function ‘phyper’ in R v.4.3.1 (R Development Core Team, 2019).

## Results

### The impact of Azolla co-cultivation on rice metabolome

The phenotype of rice roots and aerial organs was significantly modified by co-cultivation with Azolla (Fig. 1b, c), as shown in detail in the companion paper by Cannavò et al. (2024, Preprint). In our non-targeted metabolomic analysis, we compared the metabolome of leaves and roots of rice co-cultivated with Azolla to those of plants grown without Azolla at two time points, 40 and 60 days after the onset of co-cultivation (doc). Overall, we detected 15348 and 14574 metabolite-related mass-features (MFs) (after de-isotoping and peak-grouping of clusters and adducts) in rice leaves and roots, respectively (Fig. 2a, b). Among these, 6220 (40.5%) in leaves (Fig. 2a) and 6410 (44%) in roots were found (Fig. 2b), regardless of the presence of Azolla or the rice growth stage. This represents the metabolome of rice (roots and leaves) that is insensitive to either the growth with Azolla or to the plant developmental stage. Although Azolla did not supply inorganic nitrogen to the media (Cannavò et al. 2024, Preprint), it significantly induced changes in the metabolome of whole rice plants. Specifically, we detected 1268 (8.3%) and 894 (6.2%) MFs in leaves and roots, respectively, that occurred only in plants co-cultivated with Azolla (Fig. 2a, b). In particular, the co-cultivation with Azolla enhanced, over time, the number of metabolites (457 and 637 were detected after 40 and 60 doc, respectively) in rice leaves, whereas it slightly decreased those in roots (337 and 328 at 40 and 60 doc, respectively) (Fig. 2a, b). Besides, in rice plants grown without Azolla, 2494 (16.3%) MFs were found to be regulated in leaves (Fig. 2a) and 3005 (20.6%) in roots (Fig. 2b). In the opposite way to what happened in rice co-cultivated with Azolla, the regulation of metabolites decreased (1805 at 40 doc; 320 at 60 doc) and increased (293 at 40 doc; 2522 at 60 doc) with aging in the leaves and roots of rice grown without Azolla, respectively (Fig. 2a, b). The rice metabolome underwent, as a whole, a lower degree of regulation in plants grown with-than without Azolla, both at leaf-and root-level.

**Figure 2.**
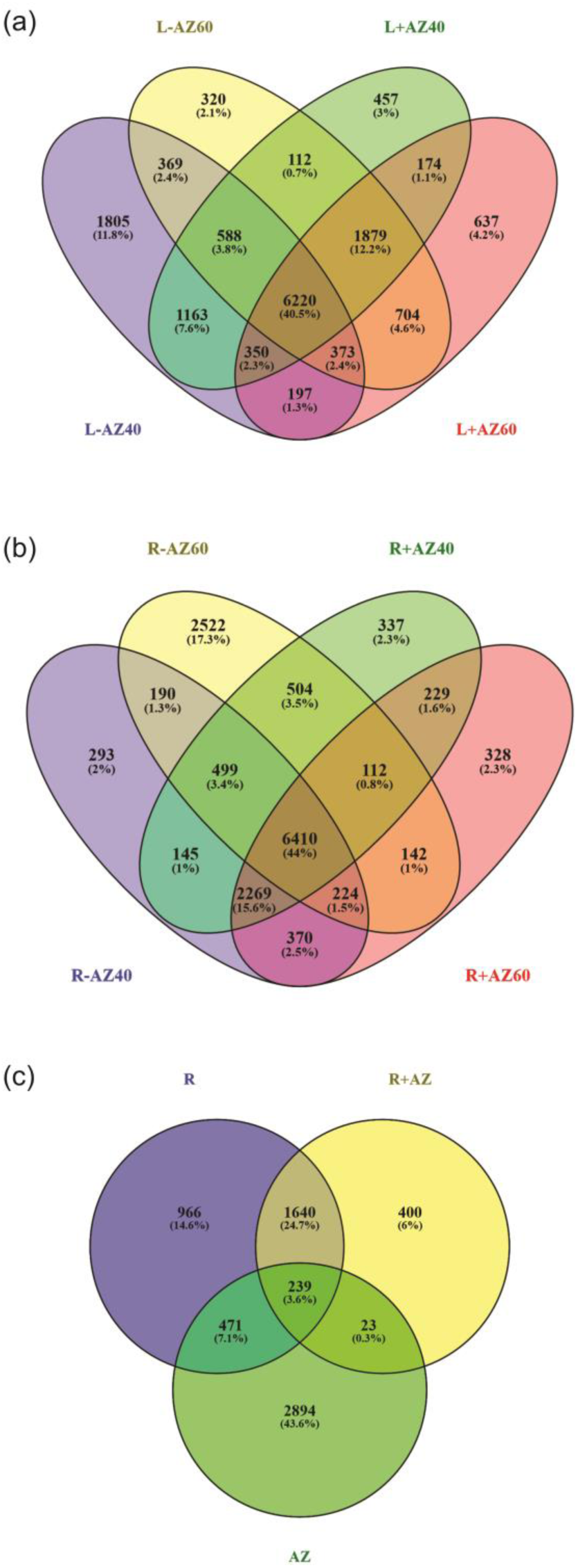
Venn diagrams showing metabolites related mass features up-and -downregulated in the metabolome of (a) leaf (L), and (b) root (R) of rice plants when co-cultivated alone (-AZ) or with Azolla (+AZ); samples were collected after 40 and 60 days of cocultivation (doc). (c) Venn diagram showing metabolites related mass features found in the culture media where Azolla (AZ), rice plants (R) and rice plants together with Azolla (R+AZ) were grown, after background correction of the respective culture solution (Watanabe for Azolla, Yoshida for rice cultivated alone or with Azolla).

We separated the effects of Azolla co-cultivation on the rice metabolome from those dependent on plant aging by using the multivariate statistical approach OPLS-DA (Fig. 3a, b). This analysis clearly showed significant differences in the metabolomes of both leaves and roots of rice due to Azolla co-cultivation at both 40 and 60 doc (Fig. 3a, b). In leaves, the number of upregulated metabolites was higher than those downregulated at both plant growth stages (Table 1). The impact of Azolla co-cultivation on rice leaf metabolome increased over time, resulting in a higher number of both upregulation and downregulation of metabolites at 60 doc compared to 40 doc (Table 2; Fig. 4 -upregulated; Fig. S1 -downregulated). Among these metabolites, most (∼ 70%) showed a low degree of regulation (Log2FC between 0 and 1) at 40 and 60 doc. In addition, while the number of highly upregulated metabolites (Log2FC >3) decreased over time, the number of those highly downregulated (Log2FC <3) increased from 40 to 60 doc (Table 1).

**Figure 3.**
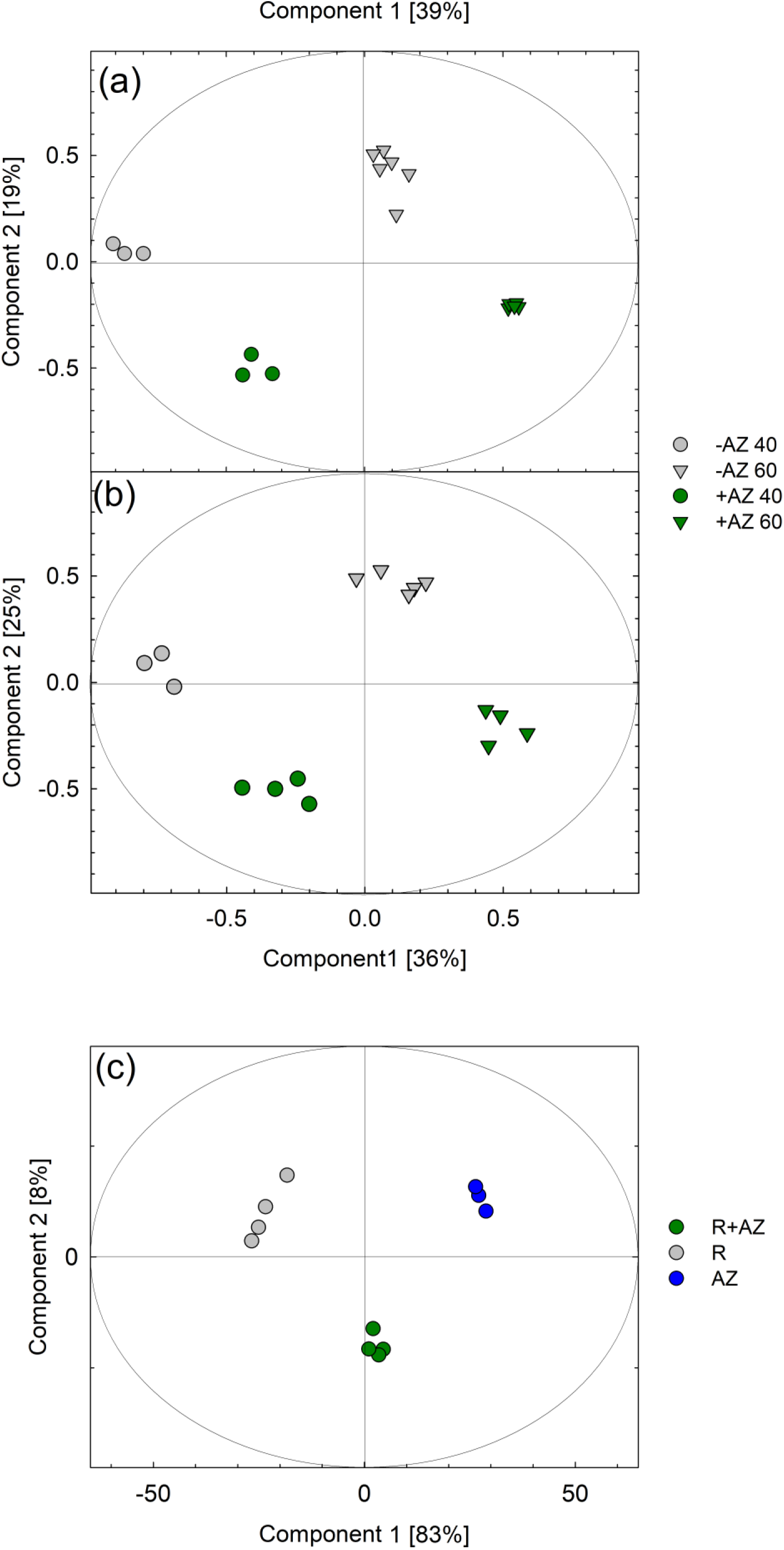
Score plots of orthogonal partial least square regression discriminant analyses (OPLS-DA) showing the variance of metabolites related mass features in (a) rice leaves and (b) rice roots of plants co-cultivated with Azolla (green colour; +AZ), and without Azolla (grey colour; -AZ). Samples collected at 40 days of cocultivation (doc) are depicted with circles, and at 60 doc with triangles. (c) Score plot of OPLS-DA of culture media where rice plants were cultivated with Azolla (R+AZ), without Azolla (R), and Azolla without rice plants (AZ). The explained degree of variance of each component is given in parentheses. All OPLS-DA models were statistically significant: (a) p-value = 0.0025; (b) p-value = 0.0011; (c) p-value = 0.0004.

**Figure 4.**
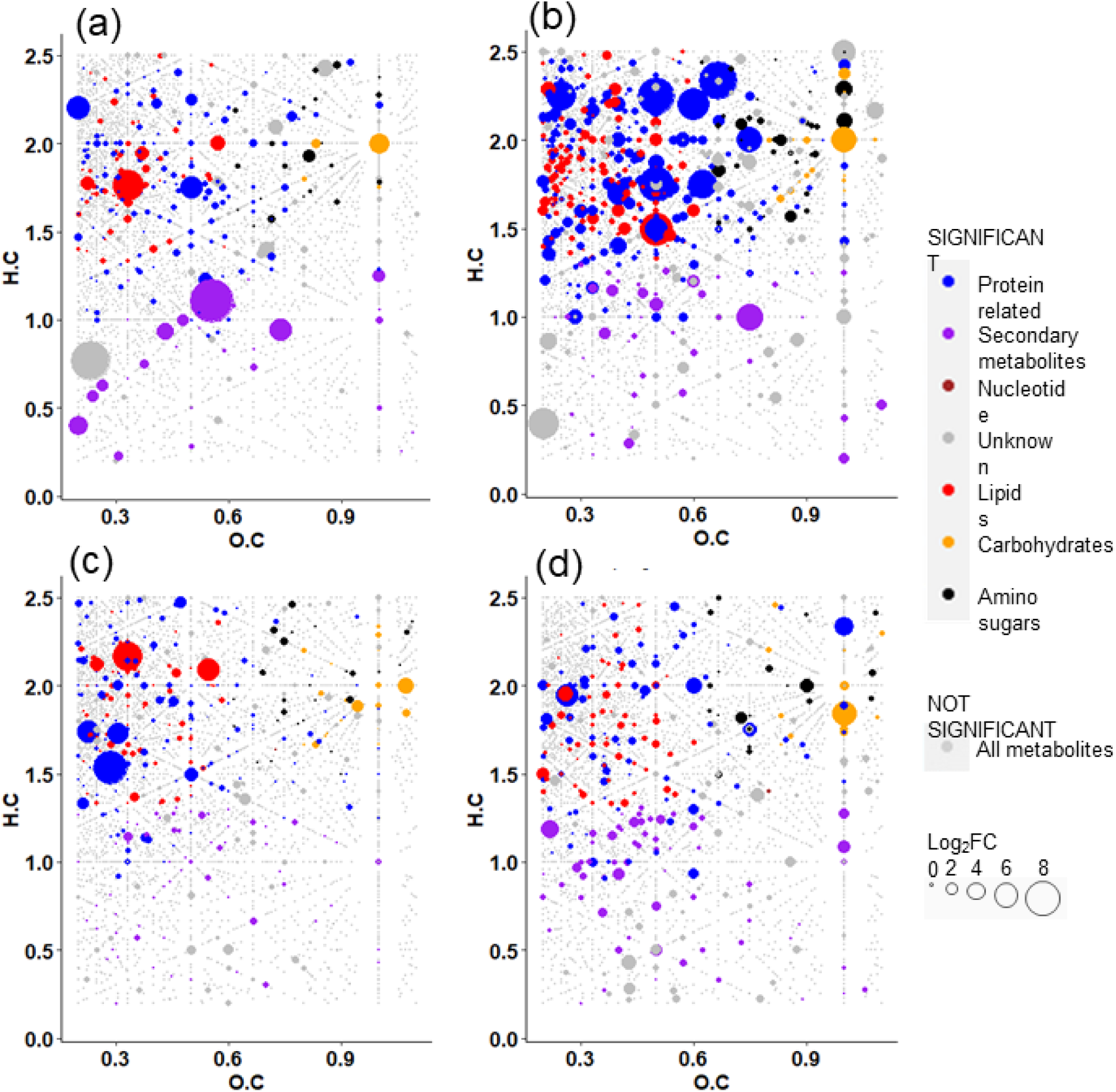
Van Krevelen diagrams of (a, c) leaf material (L) at (a) 40 and (c) 60 days; (b, d) root material (R) at (b) 40 and (d) 60 days showing significant (in colour) upregulated metabolites in presence of Azolla. According to assigned chemical formulas, the Van Krevelen diagram combined with MSCC classifies the formula-annotated mass features and assigns them to matched groups. OPLS-DA, (VIP > 1.0). In grey, not significant mass features (p<0.05, 2-way ANOVA, Benjamini-Hockberg corrected). The size of the dots reflects the log fold-change ratios between treatment (+AZ) and control (-AZ).

**Table 1.** Mass features recorded in rice plants co-cultivated with Azolla compared to those cultivated without Azolla and having significance p < 0.05 (see Fig. 4 and Fig. S1).

|  | Up-regulated |  |  |  | Down-regulated |  |  |  |
| --- | --- | --- | --- | --- | --- | --- | --- | --- |
|  | Log<sub>2</sub>fold change |  |  |  | Log<sub>2</sub>fold change |  |  |  |
|  | 0 < 1 | 1 < 3 | > 3 | Tot | -1 < 0 | -1 < - 3 | < -3 | Tot |
| Leaves 40 doc |  |  |  |  |  |  |  |  |
| Protein related | 136 | 43 | 2 | 181 | 71 | 25 | 1 | 97 |
| Lipids | 89 | 26 | 2 | 117 | 58 | 20 | ... | 78 |
| Secondary metabolites | 28 | 13 | 4 | 45 | 33 | 18 | 3 | 54 |
| Amino sugars | 21 | 9 | ... | 30 | 9 | 4 | ... | 13 |
| Carbohydrates | 13 | 2 | 2 | 17 | 10 | 3 | ... | 13 |
| Nucleotides | 2 | 2 | 1 | 5 | 2 | 1 | ... | 3 |
| Unknown | 132 | 77 | 3 | 212 | 123 | 71 | 1 | 195 |
| **Tot** | **421** | **172** | **14** | **607** | **306** | **142** | **5** | **453** |
| Leaves 60 doc |  |  |  |  |  |  |  |  |
| Protein related | 121 | 36 | 3 | 160 | 122 | 42 | 3 | 167 |
| Lipids | 84 | 23 | 2 | 109 | 65 | 20 | 7 | 92 |
| Secondary metabolites | 63 | 14 | ... | 77 | 63 | 7 | 1 | 71 |
| Amino sugars | 24 | 9 | ... | 33 | 30 | 8 | ... | 38 |
| Carbohydrates | 26 | 8 | 1 | 35 | 20 | 6 | ... | 26 |
| Nucleotides | 1 | ... | ... | 1 | 3 | 1 | ... | 4 |
| Unknown | 270 | 61 | 1 | 332 | 231 | 80 | 3 | 314 |
| **Tot** | **589** | **151** | **7** | **747** | **534** | **164** | **14** | **712** |
| Roots 40 doc |  |  |  |  |  |  |  |  |
| Protein related | 132 | 155 | 19 | 306 | 74 | 89 | 12 | 175 |
| Lipids | 152 | 148 | 4 | 304 | 72 | 45 | 6 | 123 |
| Secondary metabolites | 42 | 30 | 1 | 73 | 55 | 32 | 6 | 93 |
| Amino sugars | 23 | 21 | 2 | 46 | 22 | 16 | 1 | 39 |
| Carbohydrates | 29 | 13 | 2 | 44 | 9 | 6 | ... | 15 |
| Nucleotides | 1 | 1 | ... | 2 | 3 | 2 | ... | 5 |
| Unknown | 199 | 223 | 9 | 431 | 177 | 111 | 12 | 300 |
| **Tot** | **578** | **591** | **37** | **1206** | **412** | **301** | **37** | **750** |
| Roots 60 doc |  |  |  |  |  |  |  |  |
| Protein related | 51 | 57 | 4 | 112 | 239 | 250 | 7 | 496 |
| Lipids | 55 | 45 | 1 | 101 | 90 | 87 | 4 | 181 |
| Secondary metabolites | 40 | 39 | 2 | *81* | 46 | 31 | 5 | 82 |
| Amino sugars | 14 | 12 | ... | *26* | 35 | *27* | 1 | *63* |
| Carbohydrates | 13 | 7 | 1 | *21* | 20 | 21 | ... | *41* |
| Nucleotides | 2 | ... | ... | *2* | 2 | 1 | ... | *3* |
| Unknown | 120 | 96 | 1 | *217* | 251 | 283 | 7 | *541* |
| Tot | 295 | 256 | 9 | **560** | 683 | 700 | 24 | **1407** |

**Table 2.**
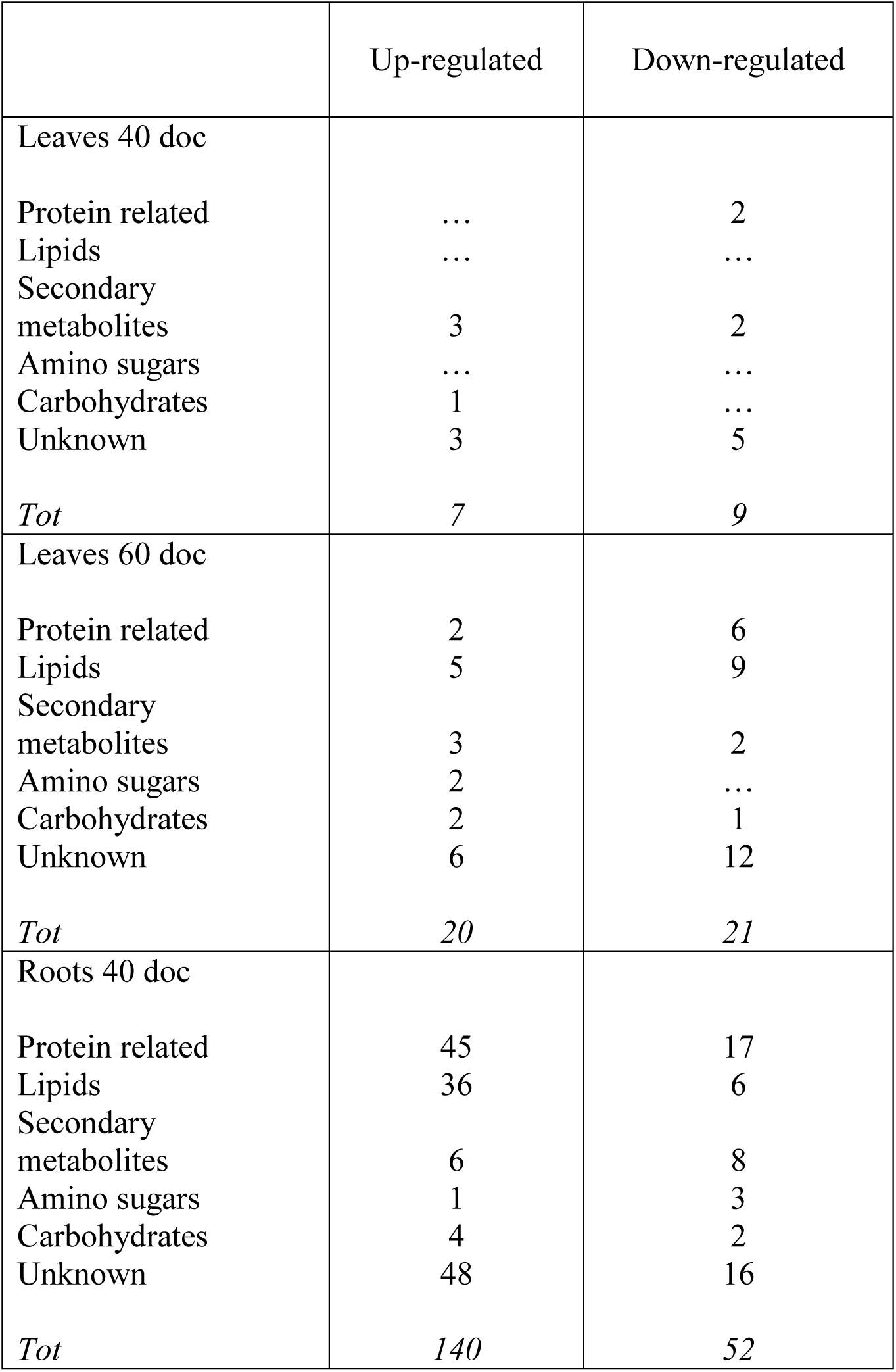
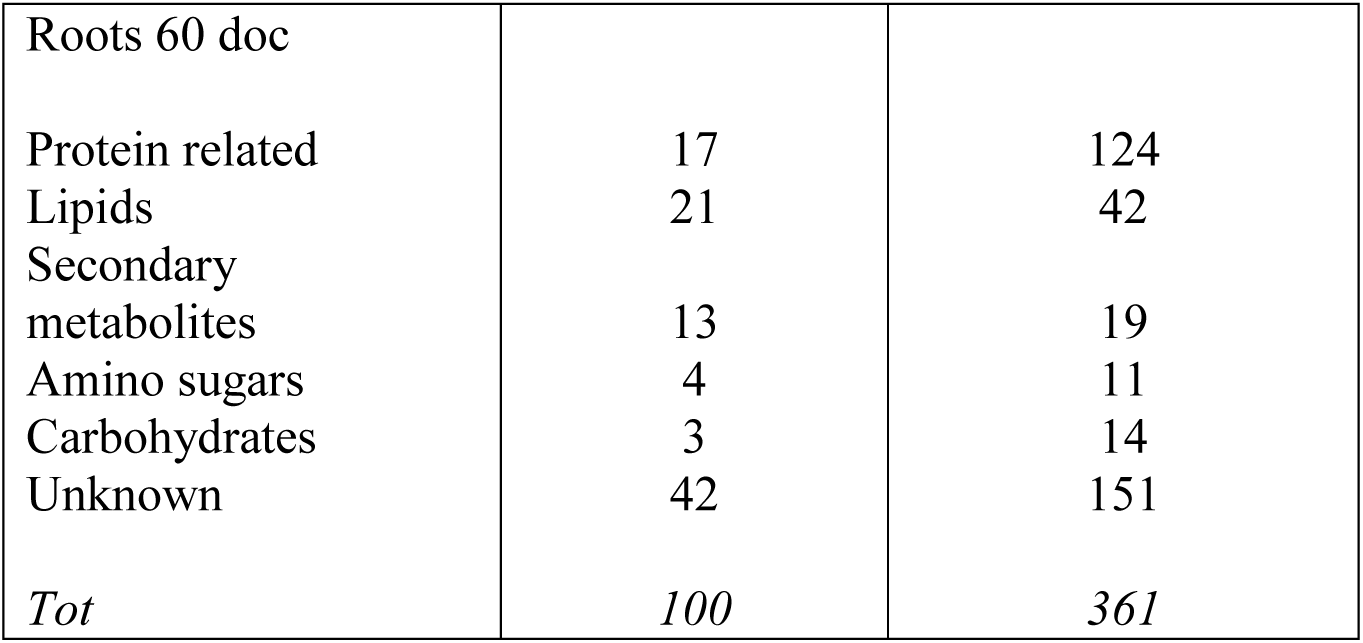
Number of strongly affected metabolites in rice plants when co-cultivated with Azolla compared to those grown without Azolla. The accounted metabolites resulted in highly discriminated (VIP > 2; OPLS-DA) and significantly changed (adjusted p-values < 0.05; Benjamin Hochberg correction) (see Table S5).

A stronger effect on metabolic regulation was found in roots than in leaves. In roots, more than 50% of metabolites increased their level up to double (Log2FC between 0 and 1) in the presence of Azolla (Table 1). Specifically, in roots of rice co-cultivated with Azolla, a higher upregulation occurred after 40 doc for metabolites with Log2fold change values between 0 and 3 (Table 1; Fig. 4), while a higher downregulation after 60 doc involved those metabolites showing a Log_2_fold change between -3 and 0) (Table 1; Fig. 4).

We classified the metabolites according to their elemental composition using the multidimensional stoichiometric compound classification (MSCC) method. This grouped compounds into broad classes such as carbohydrates, lipids, protein-related compounds (e.g., amino acids and small peptides), secondary metabolites, amino sugars, and nucleotides. We focused on significant changes (FDR < 0.05) of metabolites strongly associated with Azolla-rice co-cultivation (VIP >2, OPLS; p < 0.01, CV-ANOVA). Following this approach, we observed a higher number of metabolites changed in rice roots than in leaves (140 vs 7, at 40 doc; 100 vs 20, at 60 doc), with a plant organ-dependent shift between 40 and 60 doc. In fact, in rice roots the metabolome was upregulated at 40 doc, while in leaves it was downregulated at 40 doc compared to 60 doc (Table 2). Specifically, after 40 doc with Azolla, several protein-related metabolites (45; e.g., aminobutyric acid, leucine, dimethylarginine, alanyl-glutamine, alanyl-proline, valine-asparagine, methionine sulfoxide, 1-aminocyclopropane-1-carboxylic acid) and lipid-related metabolites (36; e.g., crotonic acid) increased in the rice root metabolome, whereas after 60 doc, the abundances of protein-related metabolites (124) and lipid-related metabolites (42) decreased (Table 1; Table S5). Interestingly, among the 36 lipid-related metabolites upregulated at 40 doc in rice roots, 7 were also upregulated at 60 doc (FDR < 0.05) including linoleic acid (Table S6). Consistent with these results, transcriptomic analysis of rice roots at 15 doc with Azolla indicated strong changes in the regulation of genes involved in amino acid salvage, i.e. processes leading to the production of amino acids from derivatives (i.e., small peptides) without *de novo* synthesis (Table 3) and transport of small peptide. In particular, the expression of the six proton-dependent oligopeptide transporter family protein, namely proton-dependent peptide (PTR) transporters, and four ATP synthase (ATP)-binding cassette (ABC) transporters, were strongly affected (Table 3). The expressions of two aminotransferases, essential to produce amino acids de novo, were also strongly downregulated (Table 3). With respect to lipids, several (14) DEGs related to the biosynthesis/metabolism of fatty acids and their transports were also differentially regulated in the roots of Azolla-cultivated rice plants at 15 doc, of which 11 were upregulated and 3 downregulated (Table S7).

**Table 3.**
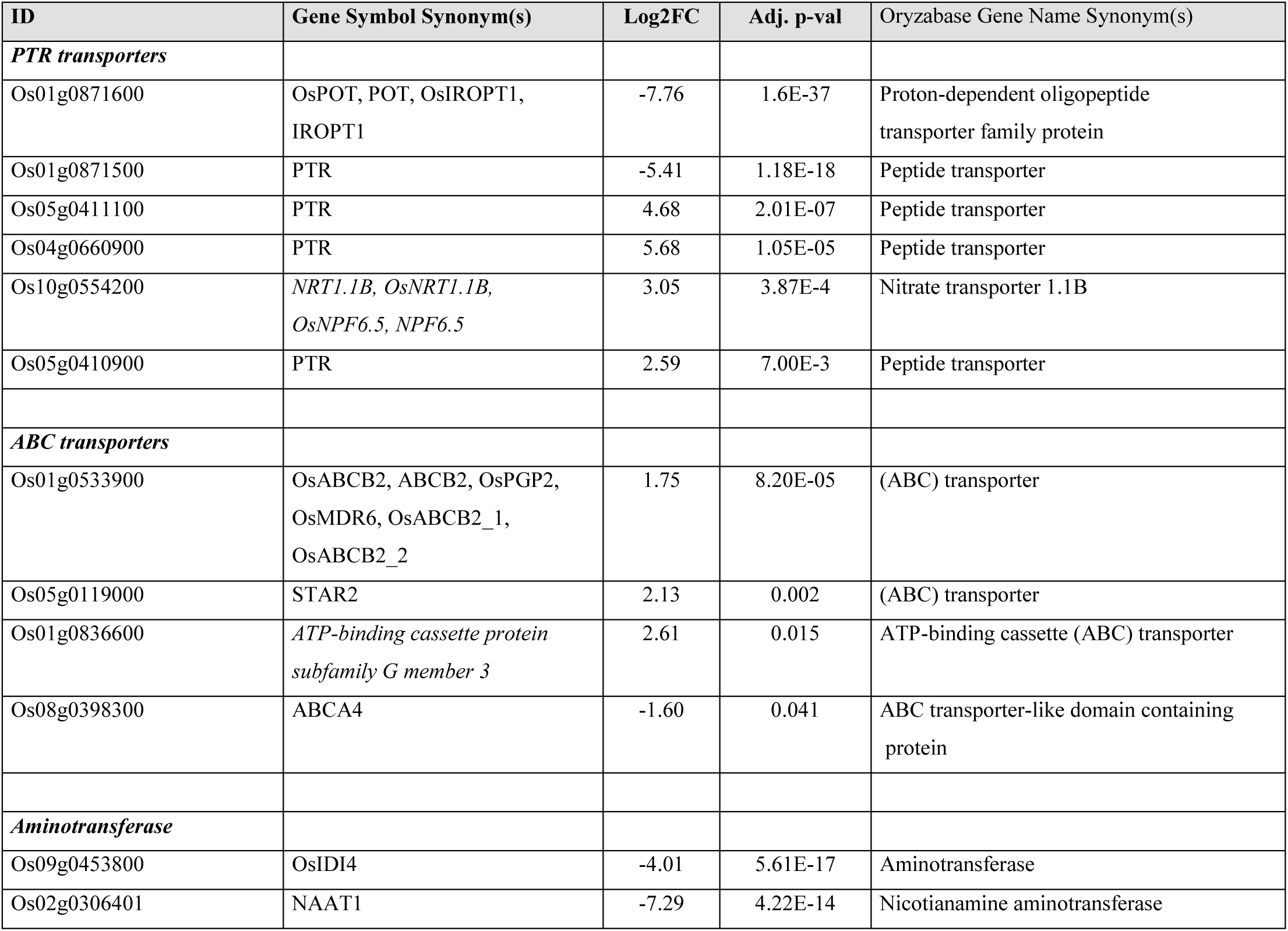
Transporter and aminotransferase-related DEGs in roots at 15 doc. The high levels of oligopeptides in the aqueous solution containing Azolla and in the roots of rice plants co-cultivated with Azolla were associated with the upregulation of several genes encoding proton-dependent peptide transporters (PTR), and ATP synthase (ATP)-binding cassette (ABC) transporters, as well as downregulation of genes encoding for aminotransferase.

In parallel with the marked reduction in the levels of metabolites in rice root following 60 doc with Azolla, we observed an increase of 20 metabolites in leaves, mainly related to lipids (5), secondary metabolites (3; i.e., flavonoids and phenolics, piperonyl aldehyde) and carbohydrates (2; i.e., xylulose-5-phosphate), as well as to some unknown metabolites (6) (Table 2; Table S3). The increasing number (from 7 to 20), over time, of the strongly upregulated metabolites in the leaves of rice co-cultivated with Azolla, may suggest that changes in rice metabolome at leaf level occurred later than those at root level following Azolla co-cultivation. Overrepresentation analysis pointed to significant up-regulation of the protein-related metabolism in leaves at 40 doc (p < 0.01, hypergeometric test) and carbohydrates at 60 doc (p < 0.01), whereas in roots of lipids at 40 doc (p < 0.001) and secondary metabolites at 60 doc (p < 0.001).

It is worth noting that many significantly regulated metabolites could not be assigned to any of the considered chemical classes by MSCC, and therefore these were referred to as ’unknown’ (Table 2; Table S5). However, we employed molecular networking (MN), a technique that can organize and visualize the chemical space in tandem mass spectrometry (MS2) data, to associate the fragmentation patterns of molecules, i.e., their chemical characteristics, with those that could be annotated through metabolomics databases. Thus, we used MN to link the ‘unknown’ metabolome to annotated metabolites present in databases. The results of this computational approach highlighted that some of the unknown metabolites whose levels increased at 40 doc were strongly associated to dipeptides (Fig. 5), supporting the observation that Azolla induces the upregulation of nitrogen metabolism in rice roots. Among the annotated metabolites whose levels strongly increase in rice leaves after 60 doc with Azolla, we found a few secondary metabolites (3) and one carbohydrate (1), as well as several metabolites related to flavonoid glycoside metabolism (Fig. 6).

**Figure 5.**
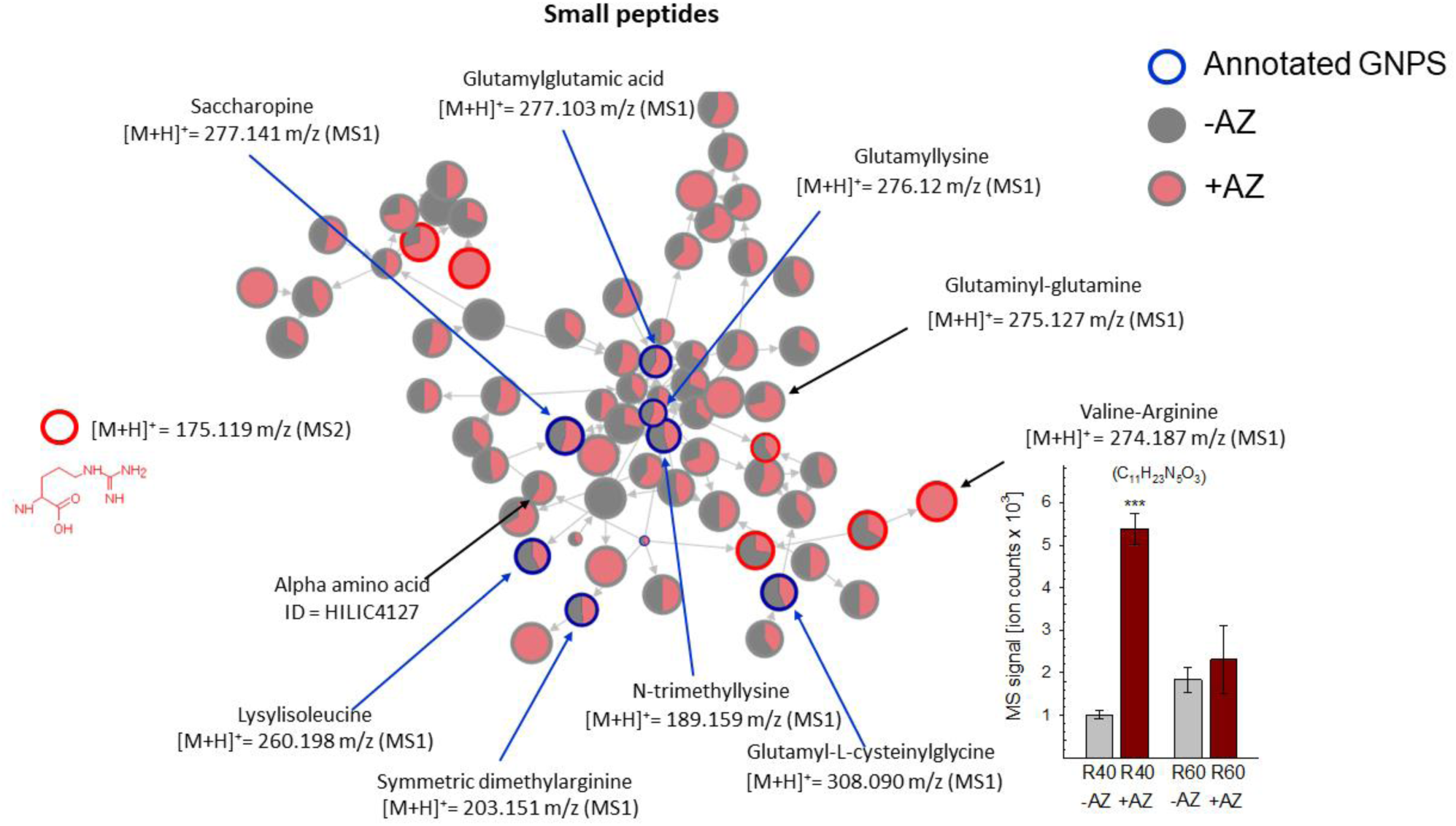
Molecular networking (MN) showing the upregulation of small peptides in roots of rice co-cultivated for 40 days with Azolla. In the network, nodes (circles) are metabolites connected via edges (nodes) based on the similarity of their mass fragmentation. The pies depict the proportion of the metabolite abundances found in rice plants co-cultivated with-(+AZ, in red) or without-(-AZ, in grey) Azolla. The blue nodes and their respective protonated ionized masses ([M+H]^+^) indicate mass features annotated as dipeptides; grey notes are unannotated mass features related to small peptides. The node sizes are the precursor intensities. MN was computed with the LC-MS data measured in HILIC(+).

**Figure 6.**
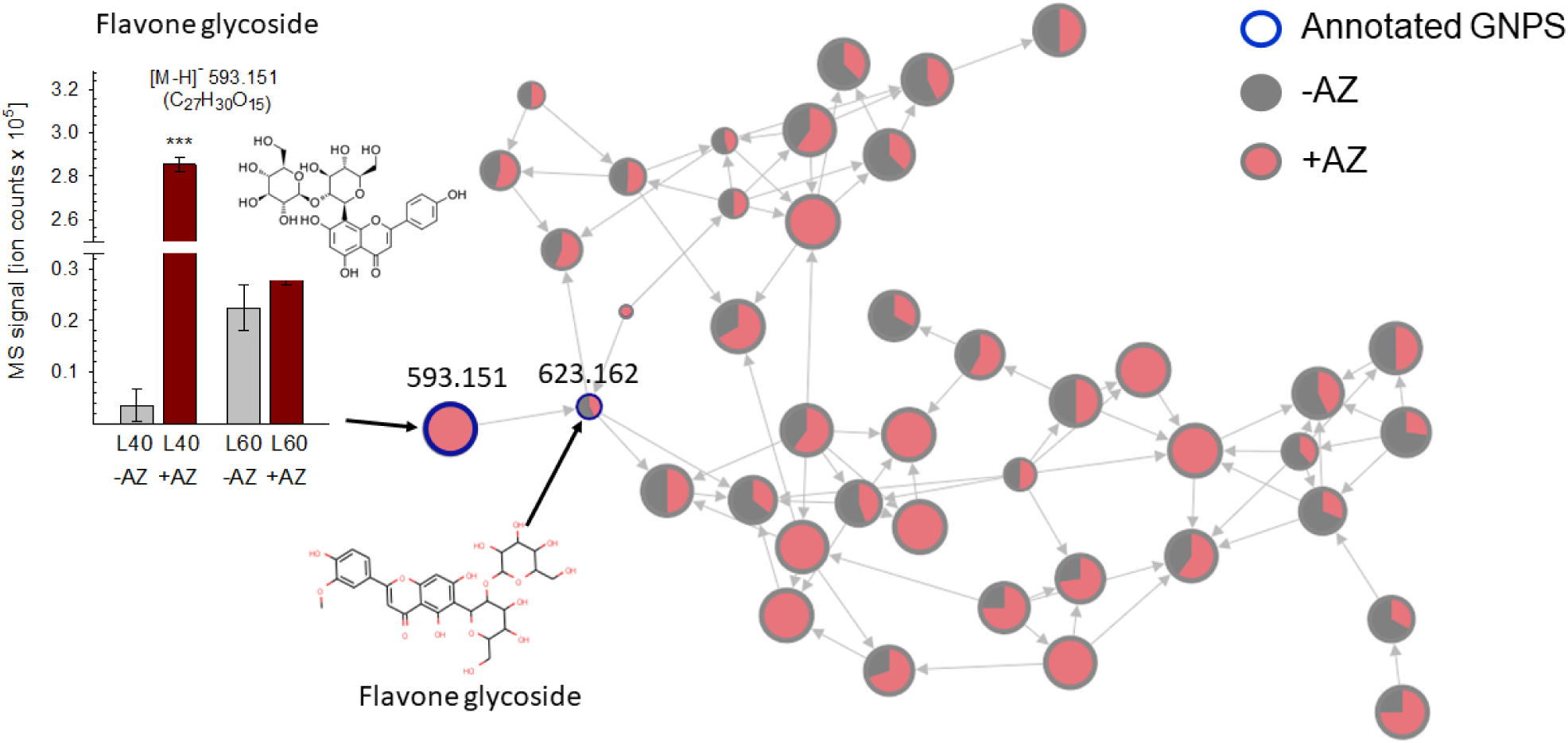
Molecular networking (MN) showing the upregulation of the flavonoid metabolism in leaves of rice co-cultivated for 40 days with Azolla. In the network, nodes (circles) are metabolites connected via edges (nodes) based on the similarity of their mass fragmentation. The pies depict the proportion of the metabolite abundances found in rice plants cultivated with-(+AZ, in red) or without-(-AZ, in grey) Azolla. The blue nodes and their respective deprotonated ionized masses ([M-H]^-^) indicate mass features annotated as flavone glycosides; grey notes are unannotated mass features related to flavonoid metabolisms. The node sizes are the precursor intensities. MN was computed with the LC-MS data measured in RP(-).

### Azolla released protein-related and flavonoid compounds in the culture medium

We analyzed the chemical compositions of the aqueous solution in which the rice plants, Azolla, and rice co-cultivated with Azolla were grown, after subtracting the compounds present in the original culture media (Watanabe and Yoshida) without plants. We detected a large number of unique metabolites (2894 MFs, 43.6%) into the culture medium when Azolla was cultivated alone (Fig. 2c). In comparison, 966 (14.6%) and 400 (6%) were the MFs only present in the medium hosting rice and rice plants co-cultivated with Azolla, respectively. Overall, the chemical compositions strongly differed, as shown by OPLS-DA analysis (p < 0.001; CV-ANOVA) (Fig. 3c; Fig. S1) possibly reflecting the effect of plant growth on two different culture media.

We, therefore, focused our analysis on the molecules released by Azolla in the culture media by comparing the samples growing in the same solution. When comparing rice with Azolla with rice alone in the same Yoshiba solution, we detected 18 metabolites upregulated, of which 2 masses could be annotated to tri-peptides. The putative release of small peptides from Azolla to the culture media was further investigated by comparing the solutions in which Azolla was grown (in Watanabe) to fresh Watanabe solution. We detected 176 MFs strongly associated with Azolla (Log2FC > 2; VIP > 1.5, p < 0.001, CV-ANOVA; Table S5). However, the library search and the MSCC approach were able to identify and classify only a few of these MF metabolites, specifically in protein-related compounds (8), amino sugar (1), lipid (8), secondary metabolite (4), and carbohydrate (2) (Table S5). Some examples are the amino acids leucine (Log2FC > 10; VIP = 17), methionine sulfoxide (Log2FC = 2.96; VIP = 1.8), phenylalanine (FC = 2.59; VIP = 1.8); 4-aminobutanoic acid (Log2FC = 4.4; VIP = 1.79); the glycerophospholipid lysophosphatidylcholine (Log2FC = 2.16; VIP = 1.77) and the fatty amide erucamide (Log2FC = 3.66; VIP = 2.02). However, by using molecular networking analysis, we observed a strong association between the remaining significant MFs and small peptides (glutamyl-cysteine, Arg-Ile, Asp-Lys, Lys-Gly-Thr) or flavonoids (e.g., quercetin-3-O-glucoside, kaempferol-3-O-glucoside, naringenin-7-O-glucoside, quercetin-3-O-manonylglucoside; Fig. 7).

**Figure 7.**
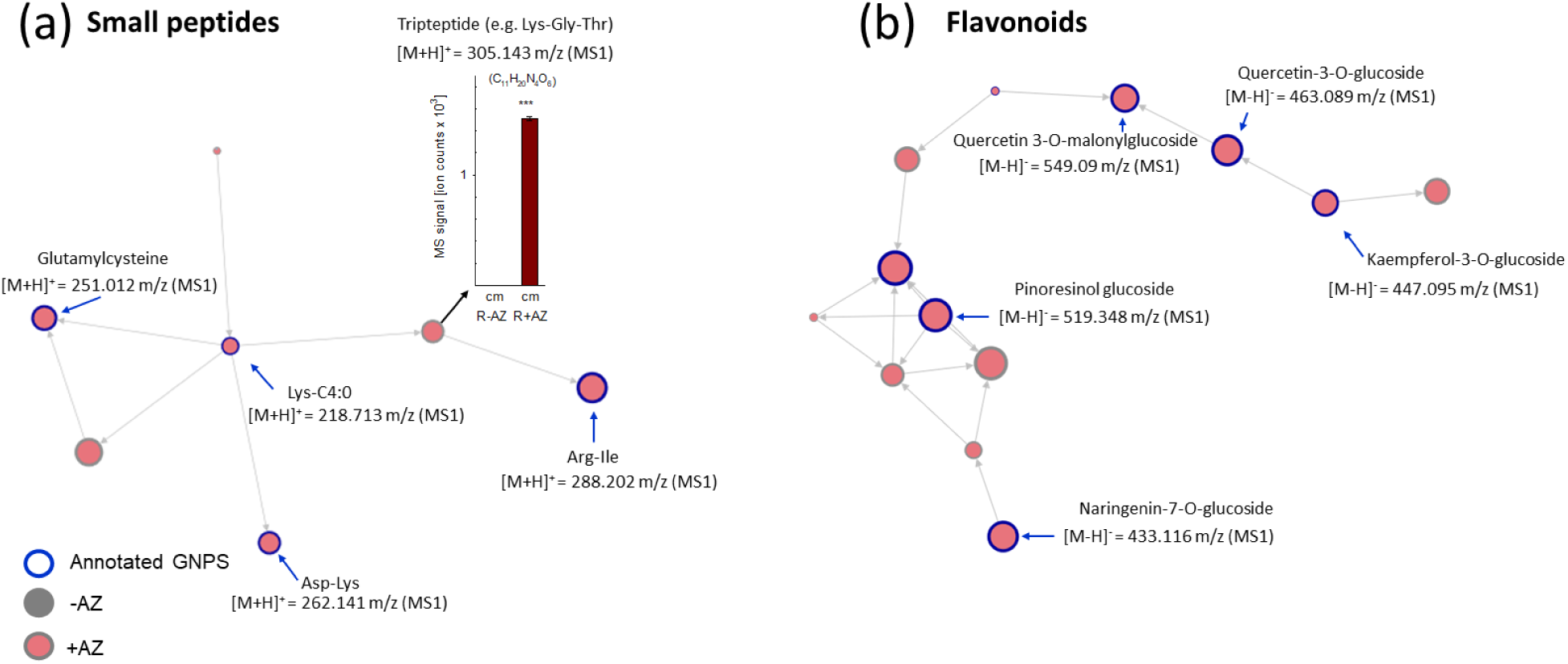
Molecular networking (MN) showing the presence of (a) small peptides (di-and tripeptides) and (b) flavonoids in the culture media of rice co-cultivated for 15 days with Azolla and compared to the medium of plants growth without Azolla. In the network, nodes (circles) are metabolites connected via edges (nodes) based on the similarity of their mass fragmentation. The pies depict the proportion of the metabolite abundances found in the medium in which rice was cultivated with-(R+AZ, in red) or without (R-AZ, in grey) Azolla. The blue nodes and their respective protonated ionized masses ([M+H]^+^ in (A) and ([M-H]^-^) in (B)) indicate mass features annotated as small peptides; grey nodes are unannotated mass features related to small peptides. The node sizes are the precursor intensities. MN was computed with the LC-MS data measured in HILIC(+) (A), and RP(-) (B).

## Discussion

In this study, we employed a non-targeted metabolomics analysis on growth media, rice roots, and leaves to deepen our understanding of the factors that induce the morphogenetic changes in rice plants as a result of the co-cultivation with Azolla (as described in the companion study by Cannavò et al. 2024, Preprint). The collected metabolic dataset was analyzed by combining the multidimensional stoichiometric compound classification (MSCC) with molecular networking (MN) to classify a large number of metabolites, including those not yet present in databases but structurally related to known compounds. Our results show that Azolla can release a wide range of phytochemicals into the aquatic culture medium that have a positive effect on rice growth and development during the early stages of their co-cultivation.

### Azolla releases metabolites into the liquid culture medium that impact the rice phenotype

The aquatic fern Azolla is known to constitutively produce volatile organic compounds (e.g., isoprene) (Brilli et al., 2022) and non-volatile metabolites (e.g., phenylpropanoids) (Costarelli et al., 2021), which can be enhanced under stressful conditions (Cannavò et al., 2023). Here, we show that Azolla can release a broad range of soluble metabolites, such as small (di-, tri-) peptides, lipids, and flavonoids, into the aquatic culture medium. These metabolites can, in turn, stimulate the growth of rice plants and influence their development (Cannavò et al. 2024, Preprint). The ability of Azolla, in association with its cyanobacterium *Trichormus azollae*, to produce metabolites may be specific to the culture medium and can also be affected by interaction with other plants. This may explain why the metabolites released by Azolla grown alone differ from those detected in the solution where Azolla was co-cultivated with rice. Nevertheless, in both solutions, we mainly detected several metabolites structurally related to protein (e.g., amino acids, di-and tripeptides), flavonoids, and lipids, indicating that Azolla and its associated microbiome can release these metabolites into the solution, regardless of the culture medium composition (Table S5). Plant roots are known to produce exudates rich in lipids, amino acids, and proteins, in addition to carbohydrates and other secondary metabolites (Canarini et al., 2019). These exudates have multiple ecological and physiological functions, such as improving plant performance (Baetz and Martinoia, 2014;) and recruiting growth-promoting rhizobacteria (PGPR) (Narasimhan et al., 2003; Upadhyay et al., 2022) by acting as signaling molecules in plant-microbe interactions (Dennis et al., 2010; Jacoby et al., 2022). Moreover, root exudates create a favorable environment for the proliferation of nitrogen-fixing symbiotic cyanobacteria, which positively impact on nutrient cycling in the ecosystem (Lu et al., 2014).

All the different classes of metabolites we found released by Azolla in the culture solution have a growth-promoting potential. The non-protein amino acids, such as exogenous aminobutyric acid, and the small peptides or free amino acids (i.e., leucine, phenylalanine, the tripeptide Lys-Gly-Thr) identified through MSCC and MN analyses in the Azolla growth medium represent a valuable source of organic nitrogen for rice that may favor stress resistance (Ma et al., 2018), and have a positive impact on root-associated bacterial communities (Wang et al., 2022a). Small peptides are known to act as hormone-like molecules in plant growth and development (Roy et al., 2018; Feng et al., 2023), function as stress signaling molecules (Chen et al., 2020) able to trigger plant defense responses (Valmas et al., 2023), and to exert beneficial effects on the microbiome of rice roots (Tejada et al., 2011; Colla et al., 2017). In addition, cyanobacterial species such as *Trichormus azollae*, are known to release free amino acids (i.e., aspartate, glutamate, alanine) into their growth medium (Thomas and Shanmugasundaram, 1992). We detected leucine and phenylalanine in the growing medium of Azolla-*Trichormus azollae*. We also identified small peptides that may serve as a source of free amino acids in the growing medium, potentially derived from larger peptides following hydrolysis-mediated by proteolytic enzymes such as proteases. These proteases may be released from Azolla or rice roots to facilitate uptake (Adamczyk et al., 2010). In turn, rice plants may take up the small peptides through transporters located on the plasma membrane (Näsholm et al., 2009; Tegeder and Masclaux-Daubresse, 2017). Consistent with this hypothesis, the gene expressions of six transporters putatively annotated as *proton-dependent peptide transporters* (*PTR*s) and four *ABC* transporters, were significantly affected in the roots of Azolla co-cultivated plants. The *PTR* gene family comprises a group of membrane transport proteins that facilitate the uptake of di-and tripeptides across cellular membranes (Stacey et al., 2002; Komarova et al., 2008). Among those 10 transporters, eight were upregulated and two were markedly downregulated in rice roots, suggesting a specific compensatory response to the high levels of di-and tripeptides present in the culture medium, e.g., by feedback inhibition of the transcription factors binding to the promoter regions of *PTR* genes. Moreover, the higher level of di-and tri-peptide in the culture medium may lower the demand for amino acid biosynthesis in rice. Accordingly, we note how the expressions of genes involved in the *de novo* aspartate and glutamate biosynthesis (i.e., *aspartate aminotransferase* and *nicotianamine aminotransferase*, respectively; Table 3) were downregulated in the roots of Azolla co-cultivated rice. We found lipids released by Azolla in the medium that might elicit plants’ innate immunity by inducing nitric oxide (NO) production, calcium influx, and oxidative burst (Erbs et al., 2003; Silipo et al., 2010; Nurnberger et al., 2004), thus sustaining plant growth under unfavorable conditions. Consistent with this, in the roots of rice co-cultivated with Azolla we detected both the differential expression of lipid transporters (Table S7), increased NO levels (see companion study by Cannavò et al. 2024, Preprint) and the presence of dimethylarginine (this study), which is produced by the methylation of arginine and is involved in NO signaling (Corpas et al., 2009; Hu et al., 2019). Likewise, two rice genes involved in (leaf) cuticular wax biosynthesis were upregulated in the roots of rice co-cultivated with Azolla. Thus, lipids and carbohydrates released by Azolla may serve to either synthesize or reinforce the external layers of rice root cells, thus providing protection against stress (Pereira et al., 2009; Ozturk and Aslim, 2010).

We also identified some flavonoids in the culture medium (Fig. 7b). Azolla is known to be rich in phenolic compounds (Brouwer et al., 2018; Costarelli et al., 2021), and here we show that it releases flavonoids into the surrounding medium (Fig. 7b). Flavonoids play a crucial role in plant stress responses by mitigating excess of reactive oxygen species (ROS) and by modulating stress signaling pathways (Daryanavard et al., 2023). Additionally, flavonoids influence phytohormone signaling, thereby contributing to the regulation of plant growth and development. In turn, rice plants may benefit from a nutrient solution enriched in flavonoids. Another compound involved in stress tolerance that we detected in the Azolla culture medium was methionine sulfoxide, which also interacts with plant growth processes (Ray and Tarrago, 2018). However, our study investigated rice growth under non-stressed conditions. Therefore, the potential ability of Azolla to increase stress tolerance of rice plants deserves further investigation under stress conditions in the future.

The higher number of metabolites we detected in the medium where Azolla was grown alone with respect to those detected in the medium from the co-cultivation (Fig. 2) suggests that the metabolites released by Azolla are taken up by the rice roots. Likewise, the lower number of metabolites detected in the medium in which rice and Azolla were co-cultivated compared to those present in the media where rice was grown alone, could be interpreted as a result of metabolite turn over due to Azolla uptake and/or chemical modifications (e.g., hydrolysis of peptides into free amino acids). Overall, since the accumulation of different forms of inorganic nitrogen was prevented in our experiments by frequent replacement of the nutrient solution, the morphogenetic changes exhibited by rice plants following co-cultivation with Azolla (see Cannavò et al. 2024, Preprint) are most likely caused by the metabolites and phytoregulators released by, and exchanged with, Azolla in the culture solution, some of which were taken up by the rice roots.

### Co-cultivation with Azolla impacts both root and leaf metabolomes

Co-cultivation with Azolla strongly affected the metabolome of rice roots and leaves. To the best of our knowledge, this is the first non-targeted metabolomics investigation of rice plants during the co-cultivation with Azolla. Previous metabolomic studies have been performed on rice plants to track the geographical origins (Hu et al., 2014; Li et al., 2022a), profile therapeutically important metabolites (Kusano et al., 2015; Rajagopalan et al., 2022), and identify biomarkers of seeds yellowing (Liu et al., 2020) and quality deterioration (Wang et al., 2022b).

In this study, we demonstrated that co-cultivation with Azolla initially affected the root-rather than the leaf metabolome, and the upregulated root metabolites were more related to the primary metabolism and, to a lesser extent, to the secondary metabolism (Fig. 4; Table 1-2). Furthermore, we observed an overall increasing number of metabolites being regulated in leaves as rice development progressed, while a decreasing number were regulated in roots, thus indicating a temporal allocation of metabolites from root to shoots. This is in good agreement with the growth-promoting effects of Azolla co-cultivation. The increased levels of metabolites in the roots of rice co-cultivated with Azolla appear to result in part from the direct uptake of metabolites released by Azolla into the culture medium (as discussed above), as well as from the activation of rice metabolic pathways triggered by signaling molecules promoted by the interaction with Azolla-released compounds.

Among the significant changes found in the rice metabolome following co-cultivation with Azolla, we observed a much higher up-regulation than down-regulation of several mass features assigned to protein-related molecules, followed by lipids, secondary metabolites, and carbohydrates. However, only a few of these metabolites showed the highest level of upregulation (Table 1). This indicates that the interaction between Azolla and rice plants is complex, involving changes in both primary and secondary metabolism similarly to those occurring between plants and growth-promoting bacteria (Mashabela et al., 2022). The systematic upregulation of metabolites in rice co-cultivated with Azolla, which initiated in roots and shifted, over time to leaves (Fig. 4), resembles the biostimulant effects shown when seaweed extracts were applied to Arabidopsis, initially affecting the roots and later producing metabolic changes in the leaves (Tran et al., 2023). We also observed a pronounced adjustment of the primary metabolism rather than the secondary metabolism in rice roots when interacting with Azolla. This effect resulted in the accumulation of protein-and lipid-related metabolites and carbohydrates (Table 2). Notably, the elevated levels of small peptides in the roots of rice co-cultivated with Azolla mirrored the enriched levels of small peptides found in the liquid medium where only Azolla was grown alone (Fig. 5; Fig. 7). This suggests that rice roots absorbed small peptides produced and released by Azolla into the medium, a finding supported by transcriptomic analysis of rice roots which revealed the strong upregulation of genes coding for small peptide transporters (Table 3).

Among the protein-related metabolites highly upregulated in rice roots during co-cultivation with Azolla, the most upregulated mass feature has been tentatively assigned to aminobutyric acid (GABA). GABA is a signaling molecule that has multiple roles in response to abiotic (Nayyar et al., 2014) and biotic (Ramputh and Bown, 1996) stresses and in modulating the plant developmental processes (Bouché and Fromm, 2004). GABA is also important for maintaining the C:N balance within the plant cells and it is involved in hormone biosynthesis and nitrogen metabolism, which is fundamental for overall plant development (Khan et al., 2021; Pei et al., 2022; Bouché and Fromm, 2004). The application of exogenous GABA to Arabidopsis has been shown to promote the direct absorption of these metabolites by the roots through the modulation of several enzymes involved in nitrogen metabolism and nitrogen uptake (Barbosa et al., 2010). In addition, exogenous GABA could impact on rice growth by affecting the roots absorption of mineral nutrients, particularly in response to the excess of iron (Zhu et al., 2020). Thus, GABA released from Azolla into the culture medium may have contributed to the changes observed in the phenotype and transcriptional profiles of iron-related genes in rice the roots (as shown in the study of Cannavò et al. 2024, Preprint).

In both rice roots and in the culture medium where Azolla was grown, we observed the accumulation of a mass feature tentatively identified as methionine sulfoxide. Methionine sulfoxide is enhanced by its rapid oxidation under increasing levels of ROS (Tarrago et al., 2015). However, under specific conditions, its biosynthesis can also occur without ROS excess, potentially through post-translational modifications (Rey and Tarrago, 2018). Methionine sulfoxide plays a crucial role in root growth, lateral root development, and root architecture regulation by interacting with auxin (Patersnak et al., 2023). Thus, methionine sulfoxide could be one of the metabolites capable of triggering the transcriptomic and morphological changes in rice plants co-cultivated with Azolla, as highlighted in in Cannavò et al. (2024, Preprint).

In the roots of rice co-cultivated with Azolla, our analysis indicates the accumulation of a mass feature that could be assigned to 1-aminocyclopropane-1-carboxylic acid (ACC). This non-protein amino acid is the precursor of ethylene, a phytohormone involved in plant development and stress response processes, which also participates in an intricate crosstalk with other hormones (Vanderstraeten and Van Der Straeten 2017). Recent studies have highlighted the role of ACC as a signaling molecule independently of ethylene, a hormone that can be transported within plants to regulate many processes including plant growth and functionality (Polko and Kieber, 2019). Thus, rice might benefit from the presence of Azolla through the stimulation of metabolites involved in phytohormone synthesis or through the direct release of hormones in the medium, as highlighted by the hormonomics analyses reported in the companion paper by Cannavò et al. (2024, Preprint).

Among the most abundant lipid-related metabolites stimulated in rice roots by co-cultivation with Azolla, we found a mass feature tentatively assigned to crotonic acid ((E)-2-butenoic acid) and another to linoleic acid. Crotonic acid has allelochemical (Jasicka-Misiaket al., 2005) and antimicrobial properties (Fang et al., 2016) and, at high levels, might improve rice competition with weeds and resistance to soil-borne diseases. Linoleic acid, whose levels increased in the roots both after 40 and 60 days of co-cultivation, is a key constituent of cellular membranes, and it is involved in the response to oxidative stress signaling (He and Ding, 2020; Saffaryazdi et al. 2020; Liang et al. 2023) and activation of defense genes (Sumayo et al. 2014). In addition, our analysis revealed an enhanced content of carbohydrates in rice roots co-cultivated with Azolla, particularly a molecule we putatively identified as glycolaldehyde. This simplest carbohydrate molecule has been detected in plant tissues (Li et al., 2022b), but its role in affecting plant metabolism is still poorly understood. Nevertheless, recent studies proposed the involvement of glycolaldehyde in the shunt pathway of photorespiration, resulting in the generation of a net gain of reducing power, which favors nitrogen assimilation and cycling within cells (Missihouna and Kotchoni 2018).

We also showed that co-cultivation with Azolla affects the metabolome of rice leaves, mainly which changes in secondary metabolites and carbohydrates (Table 1). Consistent with our study case, a metabolic reconfiguration was reported in maize leaves treated with biostimulants resulting in differential quantitative profiles of flavonoids and phenolics (Lephatsi et al., 2022). Indeed, our analysis found an increase in flavone glycoside in the leaves of rice plants co-cultivated with Azolla (Fig. 6). Flavonoid glycosides are secondary metabolites that are produced in leaves to cope with abiotic stress conditions, and their biosynthesis is stimulated during plant growth (Groenbaek et al., 2019). Since the flavonoids detected both in the roots and in the culture medium are structurally different from those we found increased in the leaves, it is likely that the accumulation of flavonoids in the leaves represents an inducible response of rice plants to the interaction with Azolla (Table S3). However, we cannot exclude that, following uptake by the roots from the culture medium, some flavonoids are translocated and undergo chemical modifications in the leaves (Buer et al., 2007). Among the flavonoids that increased in leaves of rice co-cultivated with Azolla, we found those identified as lonicerin and luteolin-7-O-rhamnoside, which are known to accumulate in response to salinity (Cai et al., 2020). Moreover, co-cultivation with Azolla induced the accumulation in the rice leaves of a mass-feature assigned to piperonyl aldehyde (piperonal). The biosynthesis of this aromatic aldehyde has been recently described in leaves of black pepper (Jin et al., 2022) and is known to play a role in plant defense against biotic stress (Bakkali et al., 2008). Thus, co-cultivation with Azolla may stimulate the synthesis of a wide array of metabolites in rice leaves, which might improve stress resistance.

Our non-targeted metabolomics analysis also detected a mass feature tentatively identified as xylulose-5-phosphate whose levels increased in the leaves of rice co-cultivated with Azolla. This five-carbon sugar is both a product and an intermediate in the pentose phosphate pathway, which is a major source of reducing power for essential non-photosynthetic processes (Kruger et al., 2003). Here, an enhanced amount of xylulose-5-phosphate in rice leaves may be the result of the induced stimulation exerted, directly or indirectly, by Azolla on rice vegetative growth, as discussed in Cannavò et al (2024, Preprint).

## Conclusions

To date, studies have primarily focused on the role of the *Azolla-Trichormus azollae* association as a nitrogen fertilizer. This study reveals that rice benefits from the co-cultivation with Azolla beyond its well-known growth promotion role as an inorganic nitrogen supplier. Our findings demonstrate that the presence of Azolla impacts the metabolome of both rice roots and leaves independently of the inorganic nitrogen levels in the medium. The modification of the rice metabolome induced by Azolla promotes growth and development within a few weeks from the onset of the co-cultivation, occurring well before the agricultural soil is enriched with inorganic nitrogen derived from Azolla decomposition.

Further investigations are required to elucidate the specific roles of the various molecules, such as small peptides and flavonoids, produced and released by Azolla, in promoting the growth (and defense) of neighboring co-cultivated crops. Nevertheless, the current study provides valuable new insights into the beneficial effects of Azolla as a biostimulant to improve rice cultivation in a sustainable and environmentally friendly way.

## Acknowledgements

We thank Marko Bertić for his technical help with the LC-MS. This work is dedicated to the memory of our eminent colleague Stefania Pasqualini (S.P.) who conceived this study, earned the grant that supported it, and co-supervised E.C. All the authors are grateful to S.P. for her substantial contribution to research and teaching activities in the field of plant physiology.

## Author contribution

FP, AG, FB: conceptualization; AG, VG, MK: methodology; EC, AG, AS, CP: formal analysis; EC, AC, SC, MC, MCV, LR, AS, CP investigation, FP, AG resources; AG, CP: data curation; FB, AG: writing -original draft; FB, AG, FP, CP, VG, MK review & editing; FB, MK, LR: funding acquisition.

## Data Availability

The data that support the findings of this study are openly available at the following link: https://osf.io/b39da/

## Supplementary data

**Figure S1** - Van Krevelen diagrams of (a, c) leaf material (L) at (a) 40 and (c) 60 days; (b, d) root material (R) at (b) 40 and (d) 60 days showing significant (in colour) downegulated metabolites in presence of Azolla. According to assigned chemical formulas, the Van Krevelen diagram combined with MSCC classifies the formula-annotated mass features and assigns them to matched groups. OPLS-DA, PLS-DA (VIP > 1.0). In grey, not significant mass features (p<0.05, 2-way ANOVA, Benjamini-Hockberg corrected). The size of the dots reflects the log fold-change ratios between treatment (+AZ) and control (-AZ).

**Table S1** - List of the macro- and micro-nuitrients in the Yoshida solution.

**Table S2** - List of the macro- and micro-nuitrients in the Watanabe solution.

**Table S3** - List of the internal standard mixture used for data normalization.

**Table S4** - Metaboscape 4.0 parameters used for processing LC-MS/MS data.

**Table S5** - Dataset of non-targeted metabolomics analysis.

**Table S6** - Mass features related to lipids shared in roots of rice grown with- and without Azolla.

**Table S7** - Lipid-related differentially expressed genes (DEGs) in rice roots sampled 15 days since the beginning of co-cultivation with Azolla. The roots of rice plants co-cultivated with Azolla affected the expression of genes involved in fatty acid metabolisms, glycerol-3-phosphate acyltransferases (GPATs), flavin adenine dinucleotide (FAD) coenzymes, and lipid transfer proteins (LTPs).

