## Supplementary figures and images for "Co-cultivation with Azolla affects the metabolome of whole rice plant beyond canonical inorganic nitrogen fertilization"

### Supplementary Figure S1

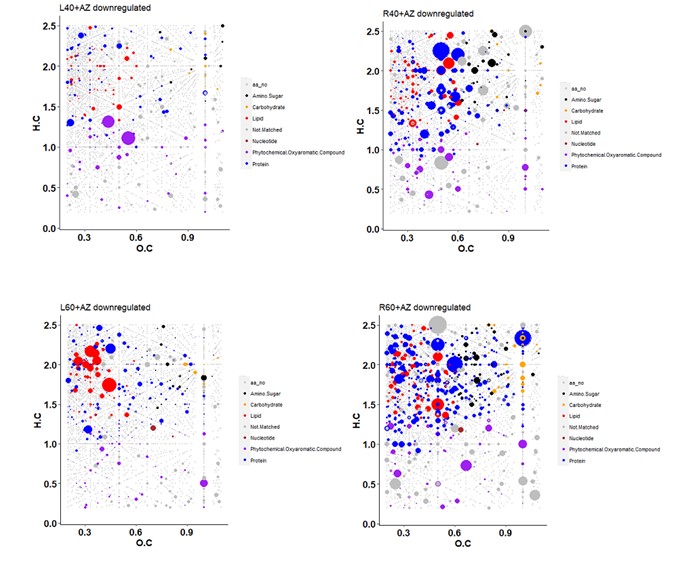
